# Modulation of Body Mass Composition using Vestibular Nerve Stimulation

**DOI:** 10.1101/087692

**Authors:** Paul D. McGeoch, Jason McKeown, Hans Peterson, V.S. Ramachandran

## Abstract

There is increasing evidence of a “set-point” for body weight in the brain, that is regulated by the hypothalamus. This system modifies feeding behavior and metabolic rate, to keep body fat within predetermined parameters. It is also known that animals subjected to chronic centrifugation show a reduction in body fat. Experiments with mutant mice found that this loss of fat appears to be mediated by a vestibulo-hypothalamic pathway. Vestibular nerve stimulation (VeNS), also known as galvanic vestibular stimulation, involves non-invasively stimulating the vestibular system by applying a small electrical current between two electrodes placed over the mastoid processes. We suggest that any means of repeatedly stimulating the otolith organs in humans would cause a reduction in total body fat, and that VeNS would be a useful technique to use in this regard. Below we provide pilot data to support this idea.

## Introduction

There is growing evidence for a “set-point” in the hypothalamus that regulates body weight, and in particular total body fat.^1–3^ This feedback control mechanism modifies feeding behavior and metabolic rate, in order to maintain body fat at a relatively fixed amount. Indeed, deviations too far in either direction from the set-point are strenuously resisted.^2,3^

Thus not only is it hard to lose weight via lifestyle changes, such as dieting, but also even if one can, maintaining the new lower weight is typically a Sisyphean task. The classic pattern being that weight will soon return back to pre-diet levels. Moreover, the set-point also explains why individuals on very high calorie diets for prolonged periods, though refraining from exercise, nonetheless gained little weight.^4^

Several nuclei within the hypothalamus are involved in regulating the set-point for body weight, but one particularly vital area appears to be the arcuate nucleus with its populations of pro-opiomelanocortin (POMC), and agouti-related peptide (AgRP) / neuropeptide Y (NPY) co-expressing neurons.^5–8^ Together these neurons comprise the central melanocortin system, which responds to levels of circulating hormones such as leptin, insulin and ghrelin, as well as nutrients such as glucose, amino acids and fatty acids.^5–8^ In the case of the POMC neurons the response is anorexigenic with a decrease in food intake and an increase in energy expenditure. Conversely, the orexigenic AgRP/ NPY neurons act in an antagonistic manner.^5–8^

The set-point for body weight obviously varies, and is thought to be determined by both genetic and epigenetic factors.^9^ It can however change over time, as illustrated by the phenomenon of middle age spread. Moreover, evidence is now emerging that Western diets with excessive quantities of certain macronutrients, in particular simple carbohydrates and saturated fatty acids damage neuronal populations within the hypothalamus and push the set-point for body fat upwards.^2,3,5^

It is also known that animals subjected to hypergravity show a marked reduction in their total body fat.^10,11^ Experiments with a mutant mouse missing the otoconia in its inner ear found that, rather than being a non-specific effect of hypergravity, this loss of fat appears to be mediated by a specific vestibulo-hypothalamic pathway.^10,11^ Here we present evidence that non-invasive electrical stimulation of the inner ear in humans appears to cause a similar loss of body fat.

## Vestibular Nerve Stimulation (VENS)

- Also called galvanic vestibular stimulation^12^
- Non-invasive technique to activate vestibular apparatus by passing current through skin overlying mastoid processes, with known good safety profile^12^
- Causes a sensation of swaying, or dysequilibrium, like being on a boat
- The experimental subjects underwent binaural AC VeNS at 0.5Hz as tolerated up to 1mA; this should preferentially activate the otoliths and sympathetic activity^13,14^
- All stimulation sessions lasted an hour, with no more than one a day

## Experiment One Methods

- Single-blind randomized controlled pilot study of pre-obese (BMI 25-30 kg/m^2^) and obese (BMI>30 kg/m^2^) subjects
- Six subjects in treatment group and 3 controls who underwent sham stimulation
- All subjects underwent dual energy X-ray absorptiometry (DXA) scanning in a fasted state at the beginning and end of study to establish body mass composition
- Realistic sham stimulation device with electrodes to skin overlying both mastoids
- All subjects underwent 40 hours in total of stimulation, except for initial 3 subjects in the experimental group (20 hours each)
- Stimulation sessions took place over on average a four month period
- Subjects did not change diet or exercise habits during study
- DXA scans gave a break down on fat distribution including truncal fat, which we took as a surrogate of visceral fat, which is particularly associated with the diseases of obesity
- Data analyzed using unpaired t-test, with two tailed significance values, and variance assumed to be equal

## Experiment One Results

The treatment group had: 3 females and 3 males; a median age of 23 years (range 21 to 64); and a median BMI of 33.8 (range 25.4 to 50.2). The control group had: 2 females and 1 male; a median age of 23 years (range 19 to 64); and a median BMI of 28.7 (range 25.3 to 34.7).

The treatment group demonstrated a significant reduction in truncal fat (t(7)=2.88, p<0.025), as well as a trend towards a significant reduction in total body fat (t(7)=2.17, p=0.067). Overall the treatment group showed an 8.3% decrease in truncal fat (cf. an 8.6% increase in the control group), and a reduction in total body fat of 6.3% (cf. a 6% increase in the controls). Note the control group was skewed by a subject who gained 3.8kg in truncal fat. However, even if the weight gained by her is discounted to zero the difference in truncal fat between groups remains significant (t(7)=2.53, p=0.039).

## Experiment Two Methods and Results

Six fasted subjects (3 with BMI 20-25; 3 with BMI>25), underwent two one hour stimulation sessions on different days. One was an hour of VeNS as described above, and the other was a sham session. Thus each subject acted as their own control. All sessions started at 8am and subjects gave saliva samples and rated the intensity of their appetite (on a linear rating scale from 0 to 10), every 30 minutes for 150 minutes.

The saliva samples were analyzed using validated ELISA techniques for leptin and insulin. The collated, smoothed results, displayed graphically in Figures 1–3 as a change from point zero at baseline, suggest VeNS acts to reduce appetite and release anorexigenic hormones (insulin and leptin) that decrease food intake and increase energy expenditure.^3,7,8^

**Figure 1.**
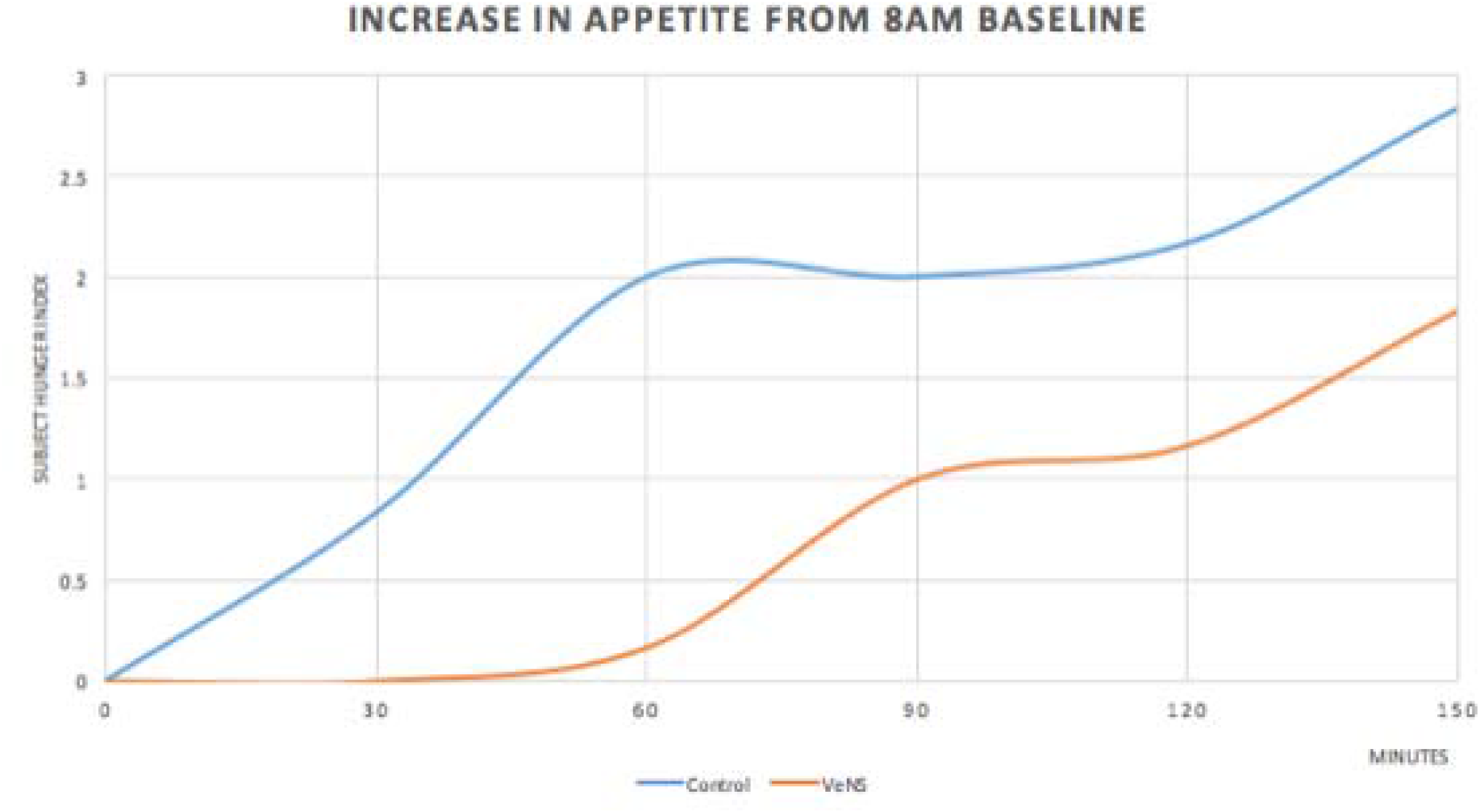
Change in appetite from baseline. Control session (blue) & VeNS session (orange). Note the reduced appetite after the one-hour VeNS session persists beyond the stimulation period.

**Figure 2.**
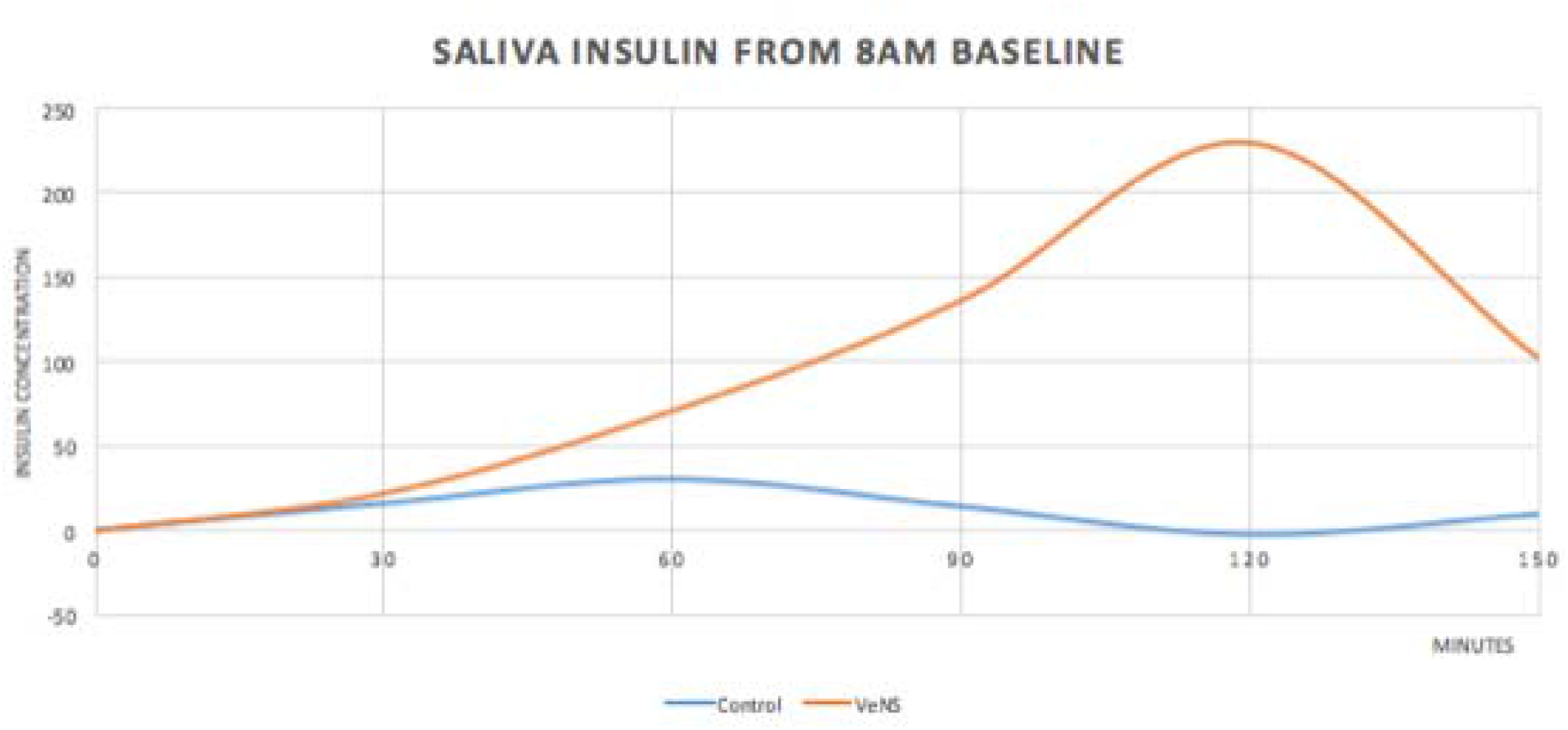
Insulin change from baseline. Control session (blue) & VeNS session (orange). Note insulin continues to rise after the one-hour VeNS session, as is seen after a meal.

**Figure 3.**
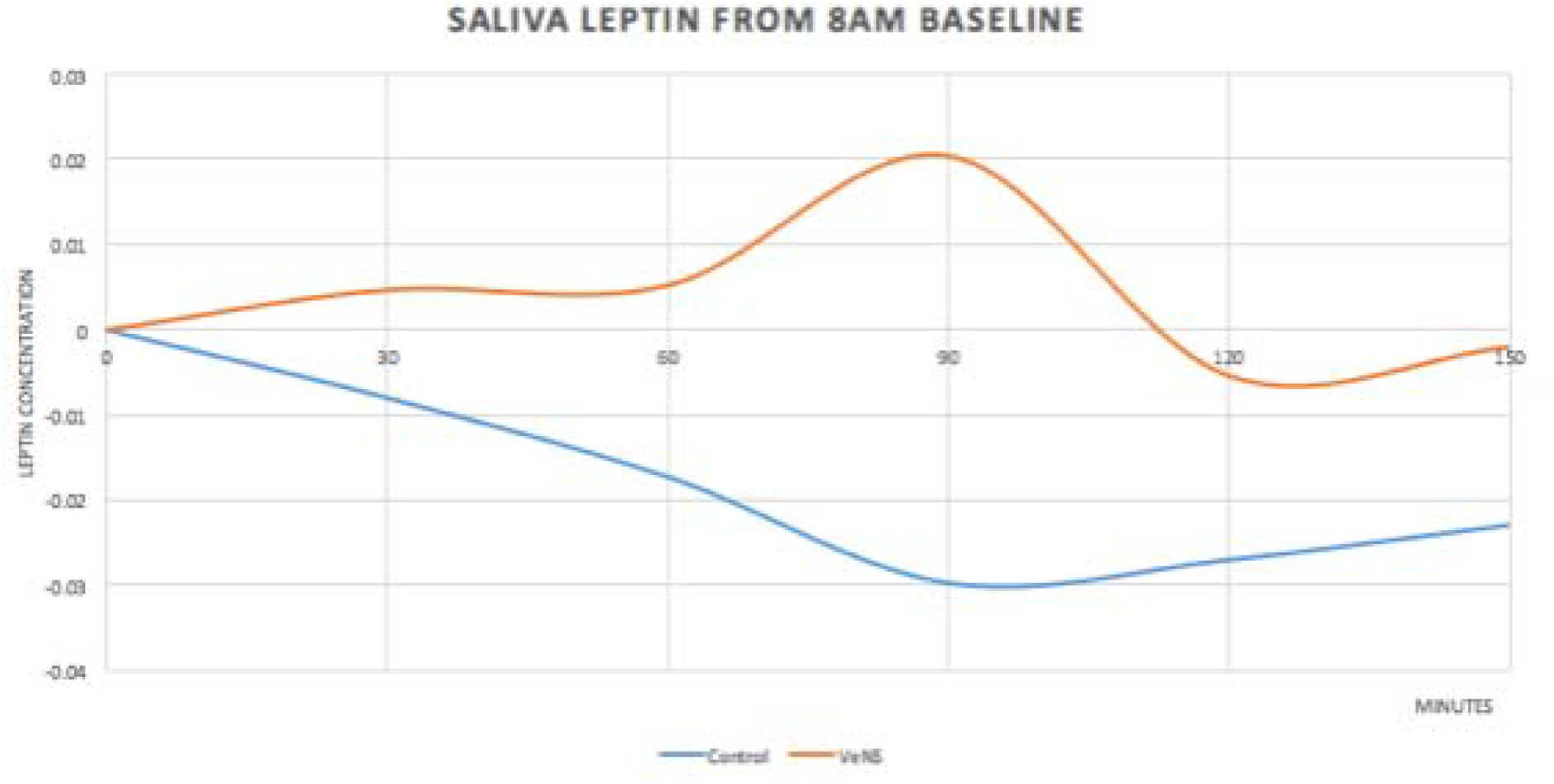
Leptin change from baseline. Control session (blue) & VeNS session (orange). Note leptin continues to rise after the one-hour VeNS session, as is seen after a meal.

## Discussion

Our data suggest that VeNS leads to a similar loss of fat in humans that vestibular stimulation is already known to produce in animals. This finding raises three questions:

1. Does the neuro-anatomical architecture exist to allow vestibular stimulation to mediate this effect?
2. Is it possible to alter the set-point for body weight?
3. Why would activating the otolith organs, which detect linear acceleration and gravity, cause a reduction in body fat?

There is growing evidence that the answer to the first two of these questions is yes.

It is recognized that the vestibular system plays an important role in homeostasis, and vestibular input is known to project to brainstem homeostatic sites, especially the parabrachial nucleus, before traveling onwards to the hypothalamus.^15,16^ Animal studies have similarly found vestibular stimulation causes robust activation of the arcuate nucleus of the hypothalamus,^11^ which sits at the crux of the central melanocortin system. And recent work has revealed melanocortinergic neurons in the medial vestibular nucleus; the main projection target of the utricle.^17,18^ Similarly, an enzyme that modulates melanocortin signaling is expressed in vestibular nuclei,^19^ and POMC neurons in the nucleus of the solitary tract, a major parasympathetic site, also receive input from the medial vestibular nucleus.^20^ All this suggests that the vestibular system has a role in the central melanocortin system, which regulates the body weight set-point.

As for whether interventions can modulate the set-point, there’s evidence polyunsaturated fatty acids may do just that, and similarly long term reduction of sugars and saturated fats in the diet may eventually lead to a downwards calibration in the set-point by allowing hypothalamic neurons to recover.^2^ It is also now increasingly believed that bariatric surgery actually works by reducing the central set-point, rather than by decreasing caloric intake.^21^ We suggest that VeNS reduces body fat by altering the hypothalamic set-point for body weight. We propose that otolith organ stimulation lowers the set-point for body fat, because otolith activation is taken by the brain to indicate a state of increased physical activity. As such it is more efficient to change to a leaner physique, in order to avoid wasting energy. This is in keeping with, indeed underlines, the fundamental role of the vestibular system in homeostasis.^15^ VeNS may thus constitute a new approach in the treatment of excess body fat and obesity.

## Declaration

PDM and VSR are named as co-inventors on a pending patent application filed by the University of California pertaining to the use of vestibular stimulation to modify body mass composition. PDM & JM have co-founded a company to license and commercially develop this technology.

## Acknowledgements

We thank Bill Rosar and J. Smythies for discussions.

